# Fertility decline in female mosquitoes is regulated by the *orco* olfactory co-receptor

**DOI:** 10.1101/2023.01.07.523099

**Authors:** Olayinka G. David, Kevin M. Sanchez, Andrea V. Arce, Andre Luis Costa-da-Silva, Anthony J. Bellantuono, Matthew DeGennaro

**Affiliations:** Department of Biological Sciences & Biomolecular Sciences Institute, Florida International University, Miami, FL 33199, USA

## Abstract

Female *Aedes aegypti* mosquitoes undergo multiple rounds of reproduction, known as gonotrophic cycles. These cycles span the period from blood meal intake to oviposition. Understanding how reproductive success is maintained across gonotrophic cycles allows for the identification of molecular targets to reduce mosquito population growth. Odorant receptor co-receptor (*orco*) encodes a conserved insect-specific transmembrane ion channel that complexes with tuning odorant receptors (ORs) to form a functional olfactory receptor. *orco* expression has been identified in the male and female mosquito germline, but its role is unclear. We report an *orco-*dependent, maternal effect reduction in fertility after the first gonotrophic cycle. This decline was rescued by repairing the *orco* mutant locus. Eggs deposited by *orco* mutant females are fertilized but the embryos reveal developmental defects, reduced hatching, and changes in ion channel signaling gene transcription. We present an unexpected role for an olfactory receptor pathway in mosquito reproduction.

**HIGHLIGHTS:**

- Loss of the *orco* olfactory co-receptor promotes female mosquito fertility decline.
- After their first reproductive cycle completes, *Ae. aegypti orco* mutant females produce embryos with disrupted development and reduced hatching.
- CRISPR/Cas9-mediated repair of the *orco* mutation rescues the fertility defect.
- Gene expression profiling of embryos from *orco* mutant females supports a role for ion channel signaling in mosquito development.

## INTRODUCTION

*Aedes aegypti* mosquitoes are competent vectors of arboviral pathogens including Zika and dengue. The global spread of *Ae. aegypti* populations is a result of their ability to colonize new climates ^1–3^. *Ae. aegypti* females produce drought-resistant eggs capable of surviving for months under laboratory conditions ^4^. Egg resistance to desiccation is thought to be conferred by an inner eggshell layer, the serosal cuticle, secreted during early embryogenesis ^5^, and gives unhatched larvae a survival advantage under harsh conditions ^6^. As global temperatures continue to rise, *Ae. aegypti* mosquitoes are predicted to expand their range, putting new regions at risk of vector-borne disease ^2,7,8^. Suppressing mosquito populations using novel, molecularly informed approaches remain a promising strategy for vector control ^9–11^. Research efforts have mainly focused on aspects of mosquito reproductive behavior leading to successful mating ^12–14^, and the subsequent physiological changes that are induced in the female ^15–17^. However, female mosquito fecundity, the potential to lay eggs, as well as fertility, the successful production of offspring, remains poorly understood.

Anautogenous mosquitoes such as *Ae. aegypti* require vertebrate blood to complete egg development ^18^. In *Ae. aegypti*, blood meal intake stimulates the release of two hormones, ecdysteroidogenic hormone (OEH) and insulin-like peptides (ILPs), from secretory cells in the brain ^19,20^. These neurohormones trigger a cascade of events leading to the release of primary egg chambers from previtellogenic arrest to complete oocyte maturation ^20–23^. Under laboratory conditions, a female *Ae. aegypti* can undergo multiple rounds of reproductive cycles, also known as gonotrophic cycles, during her lifespan ^24^. A gonotrophic cycle spans the events starting from host seeking for a blood meal to egg development, and ultimately, the deposition of mature eggs. Pathogen transmission by mosquitoes is primarily horizontal, requiring a female mosquito to first ingest an infectious blood meal from one host before transmitting it to another ^25^. However, vertical transmission of pathogens has been documented ^26^. The close association between multiple blood feeding events or gonotrophic cycles with infection makes successive reproductive cycles particularly crucial for understanding disease transmission and mosquito population dynamics.

During oogenesis, maternal transcripts are loaded into the oocyte ^27^. Many of these maternal mRNAs are maintained in the mature oocyte until they are translated post-fertilization to drive the earliest events of embryonic development ^27^. In many dipterans including *Drosophila melanogaster, Ae. aegypti,* and *Anopheles* mosquitoes, the anterior-posterior and dorsal-ventral axes of the embryo are specified by localized maternal transcripts in the oocyte ^28–31^. Classic examples of maternal effect genes in *Drosophila* include *bicoid* (*bcd*) and *nanos* (*nos*) which specify embryonic anterior and posterior poles, respectively ^31^,^32^. Both *bcd* and *nos* are localized to their respective presumptive pole in the developing oocyte by an RNA binding protein, *staufen* (*stau*), and are translated in the early embryo ^33^. These maternal factors are distributed in gradients along the embryonic axes for proper embryo patterning ^34,35^.

To transition from a specialized gamete to a totipotent embryo, a mature oocyte must first be activated. Egg activation is a conserved process that occurs in both invertebrates and vertebrates ^36–39^. In vertebrates and some invertebrates, fertilization is a prerequisite for egg activation ^40,41^. However, ovulation, not fertilization, mediates egg activation in insects ^42^. A characteristic feature of egg activation across taxa is calcium influx from extracellular sources ^38,43^. Mechanosensitive Transient Receptor Potential (TRP) channels have been identified in *Drosophila* which mediate calcium influx to generate an initial calcium wave during egg activation ^39,44,45^. Additional calcium from internal storage in the endoplasmic reticulum released through an IP_3_-mediated pathway is necessary for calcium wave propagation ^39^. In addition, phospho-regulation of several maternal proteins occurs upon egg activation ^46^. Germline knockdown of candidate maternal genes that are phospho-regulated during egg activation produces a range of phenotypes including reduced or abolished egg production and hatching ^47^. Taken together, the contribution of multiple ion channel dependent processes is required for egg-to-embryo transition and early embryogenesis.

As embryonic development progresses, maternal gene products are gradually degraded, allowing for the zygotic control of embryogenesis in a process termed the maternal-zygotic transition (MZT) ^48^. In *Drosophila*, the insect model system for which the MZT has been most thoroughly characterized, the first several synchronous nuclear divisions after fertilization are primarily driven by maternal gene products deposited in the oocyte during egg development ^49^. Gradual decay of these maternal products precedes the zygotic genome activation (ZGA) ^49,50^. ZGA has been shown to correlate with the activity of maternally encoded transcriptional regulators such as Zelda (*zld*) and Smaug (*smg*), indicative of a coupled role for maternal gene products in removing maternal transcripts and regulating zygotic genome activation during embryogenesis ^51,52^.

Odorant receptor co-receptor (*orco*) is an insect olfactory receptor that makes odor-responsive ion channel complexes with a family of ligand-selective odorant receptors (ORs) ^53^. Interestingly, *orco* and several ORs are expressed in ovaries, testes, and developing *Ae. aegypti* embryos ^54,55^. *orco* has been linked with temperature-sensitive defects in sperm release in *Drosophila* ^56^, as well as declines in the number of eggs laid in the brown planthopper, *Nilaparvata lugens* ^57^. Moreover, testicular olfactory receptors have been shown in *in vitro* assays to mediate sperm chemotaxis in humans ^58^, and sperm activation in *Ae. aegypti* and *An. gambiae* mosquitoes ^55^. However, the *in vivo* function of ORs expressed in mosquito gonads and embryos requires further investigation.

In this study, we report an *orco*-dependent maternal effect fertility decline in *Ae. aegypti*. In the second gonotrophic cycle, *orco* mutant females produce a similar number of eggs but show significantly more embryonic lethality when compared to controls. Eggs produced by *orco* mutant females in the second reproductive cycle are fertilized but most of the resulting embryos fail to hatch. These embryos showed a robust pattern of differential expression of membrane ion channel signaling mRNAs when compared to controls. A CRISPR/Cas9-mediated repair of the deletion in the *orco^16^* mutant locus was sufficient to rescue the fertility phenotype. Our results strongly suggest that *orco* plays an early role to sustain embryonic development and hatching in *Ae. aegypti* mosquitoes.

## RESULTS

### *orco* is necessary for sustained fertility over multiple gonotrophic cycles in *Ae. aegypti*

A previous transcriptomic study reported notable expression of odorant receptors (including *orco*) throughout *Ae. aegypti* embryo development ^54^. In another study, *orco*-mediated sperm activation was demonstrated in *in vitro* assays in *Ae. aegypti* ^55^, suggestive of a potential role for *orco* in mosquito reproduction. To test this hypothesis, we assayed the fecundity and fertility of a previously characterized *orco* mutant in *Ae. aegypti* ^59^, over successive gonotrophic cycles (Figures 1A & 1B). To control for recessive background mutations that may have occurred independently in either of the two *orco* mutant lines, we restricted our reproductive fitness assays to heteroallelic *orco* mutant (*orco^5/16^*) individuals generated from *orco^5/5^* and *orco^16/16^* homozygous lines. Surprisingly, we found that eggs deposited by mutant females after the first gonotrophic cycle have a significantly reduced hatching rate compared to wild-type (Figures 1C & S1A-C). Although there was no significant difference in the number of eggs produced by wild-type and *orco^516^* females in each gonotrophic cycle, females of both genotypes produced fewer eggs after the first gonotrophic cycle (Figures 1D & S1D-F). Consistent with the olfactory phenotype of this *orco* mutant ^59^, a single copy of the wild-type *orco* allele was sufficient to restore a wild-type fertility phenotype in heterozygous (*orco*^*16*/+^)individuals (Figure 2B), validating that the *orco* fertility defect is recessive.

**Figure 1.**
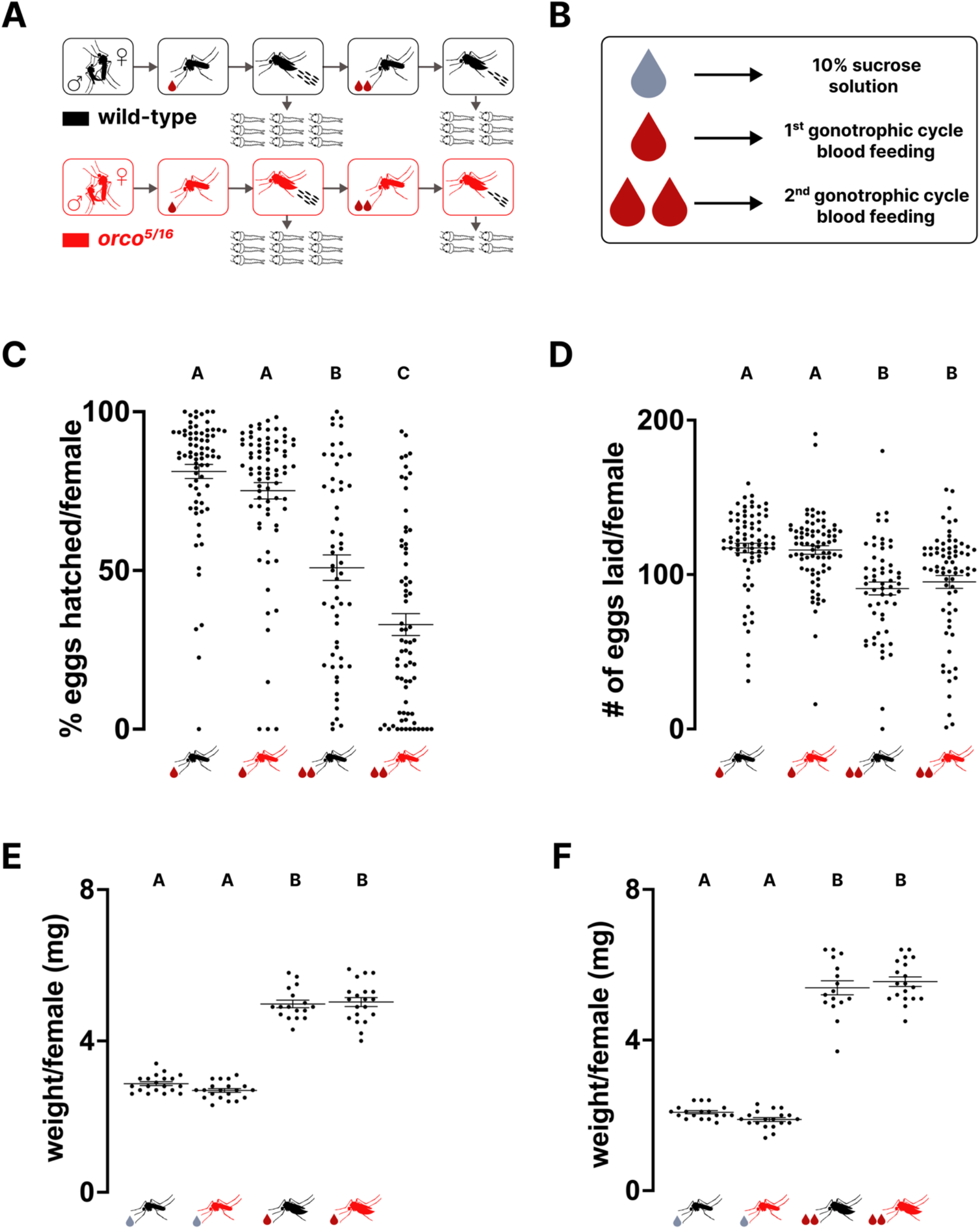
Loss of *Ae. aegypti orco* reduces fertility as the number of gonotrophic cycles increases. **(A)** Illustration of fecundity and fertility assays and phenotypes in the first and second gonotrophic cycles of *Ae. aegypti* wild-type and *orco^5/16^* females. **(B)** Symbolic representation of a female’s feeding status. Grey droplet (10% sucrose solution), single maroon droplet (first gonotrophic cycle blood meal), and double maroon droplets (second gonotrophic cycle blood meal. **(C)** Percentage of individual wild-type and *orco^5/16^* female mosquitoes eggs hatched in the first and second gonotrophic cycles (n=57-76, p<0.0001). **(D)** Number of eggs laid by individual wild-type and *orco^5/16^* female mosquitoes in the first and second gonotrophic cycles (n=56-76, p<0.0001). **(E)** Individual wild-type and *orco^5/16^* female mosquitoes weight before and after blood feeding in the first (n=17-20, p<0.0001), and **(F)** second gonotrophic cycles (n=16-19, p<0.0001). For all dot plots, long lines represent the mean, and the short lines represent standard error. Statistical analysis for C & D was done using Kruskal-Wallis test with Dunn’s multiple comparisons, and for E & F using Ordinary oneway ANOVA with Tukey’s multiple comparisons test. Genotypes marked with different letters are significantly different.

**Figure 2.**
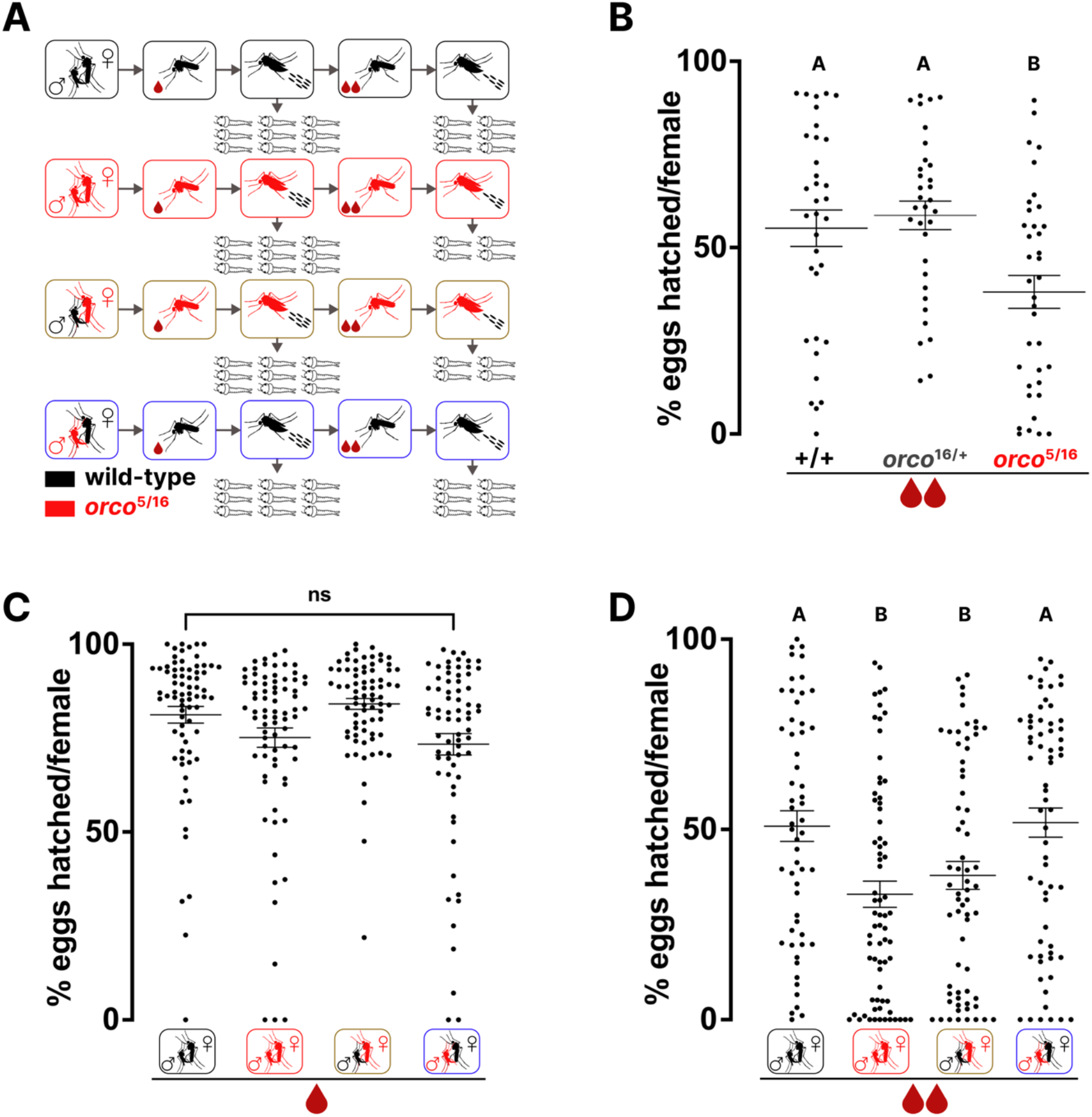
A second gonotrophic cycle maternal effect fertility decline phenotype in *orco* mutant *Ae. aegypti*. **(A)** Illustration of fecundity and fertility assays and phenotypes across 4 mating groups in the first and second gonotrophic cycles of *Ae. aegypti* wild-type and *orco^5/16^* females. **(B)** Percentage of individual wild-type, *orco^16/+^* and *orco^5/16^* female mosquitoes eggs hatched in the first and second gonotrophic cycles (n=34-37, p=0.0022). **(C)** Percentage of individual female mosquitoes of indicated genotypes & mating groups eggs hatched in the first (n=71-76, p=0.0753), and **(D)** second gonotrophic cycles (n=56-71, p=0.0005). Each virgin female mosquito was mated with a single virgin male. For all dot plots, the long line represents the mean and short lines represent standard error. All the data above were analyzed using Kruskal-Wallis test with Dunn’s multiple comparisons. Groups marked with different letters are significantly different. ns indicates no significant difference between comparisons.

Body size is a crucial factor that affects fecundity and fertility in *Ae. aegypti* females ^13,60,61^. To address whether differences in reproductive fitness resulted from differences in overall biomass, we compared the average weight of sugar-fed wild-type and *orco^5/16^* females and found no significant difference between genotypes in the first or the second gonotrophic cycles (Figures 1E and 1F). Similarly, a positive correlation is known to exist between bloodmeal size and fecundity in female mosquitoes ^62–65^. To gain insight into the average amount of blood ingested per female, wild-type and *orco^5/16^* females were individually weighed after each blood meal. We found no significant difference between genotypes in the quantity of blood ingested by females in either gonotrophic cycle (Figures 1E and 1F).

We created a mixed genotype mating assay to assess the maternal and paternal contributions of *orco* to the fertility phenotype (Figure 2A). These four mating groups are wild-type males with wild-type females, *orco^5/16^* males with *orco^5/16^* females, *orco^5/16^* males with wild-type females, and wild-type males with *orco^5/16^* females. We found no significant difference in egg hatching rate across all four mating groups in the first gonotrophic cycle (Figures 2C & S2A). We were surprised that not only the *orco^5/16^* mutant pair, but also *orco^5/16^* females crossed with wildtype males showed a reduced egg hatching rate in the second gonotrophic cycle (Figure 2D). In addition, the reduced fertility phenotype in *orco^5/16^* females remained whether virgin female and male pairs were individually mated or if mating was done in large groups (Figures 2D and S2B). Similarly, there was no significant difference between mating groups in the number of eggs produced in the first or second gonotrophic cycle (Figures S2C and S2D). Together, our results suggest that an *orco*-dependent maternal contribution is necessary for sustaining *Ae. aegypti* fertility over multiple reproductive cycles. This finding contradicts the proposed male-centric role for *orco* in mosquito reproduction ^55^.

### *orco* is required in *Ae. aegypti* females for sustained embryonic development in the second gonotrophic cycle

A female *Ae. aegypti* mosquito is inseminated by the first successful male, after which she becomes refractory to subsequent insemination ^66,67^. Indeed, a single insemination provides females of *Ae. aegypti* and *An. gambiae* with sufficient sperm for up to five gonotrophic cycles ^68,69^. Following insemination, *Ae. aegypti* females transfer sperm into a long-term storage organ, the spermathecae ^70–72^. We asked if *orco^5/16^* males successfully transfer sufficient sperm that will last for two gonotrophic cycles to their female counterparts. To test this, we quantified sperm stored in the spermathecae at two time points — prior to the first gonotrophic cycle and after the second gonotrophic cycle has been completed. The number of sperm stored by *orco^5/16^* females was not significantly different from wild-type at the two time points, showing that males of both genotypes successfully inseminate their female counterparts with an equivalent amount of sperm (Figures 3A-C). These results show that individual females of both genotypes receive sufficient sperm to fertilize her eggs over multiple gonotrophic cycles.

**Figure 3.**
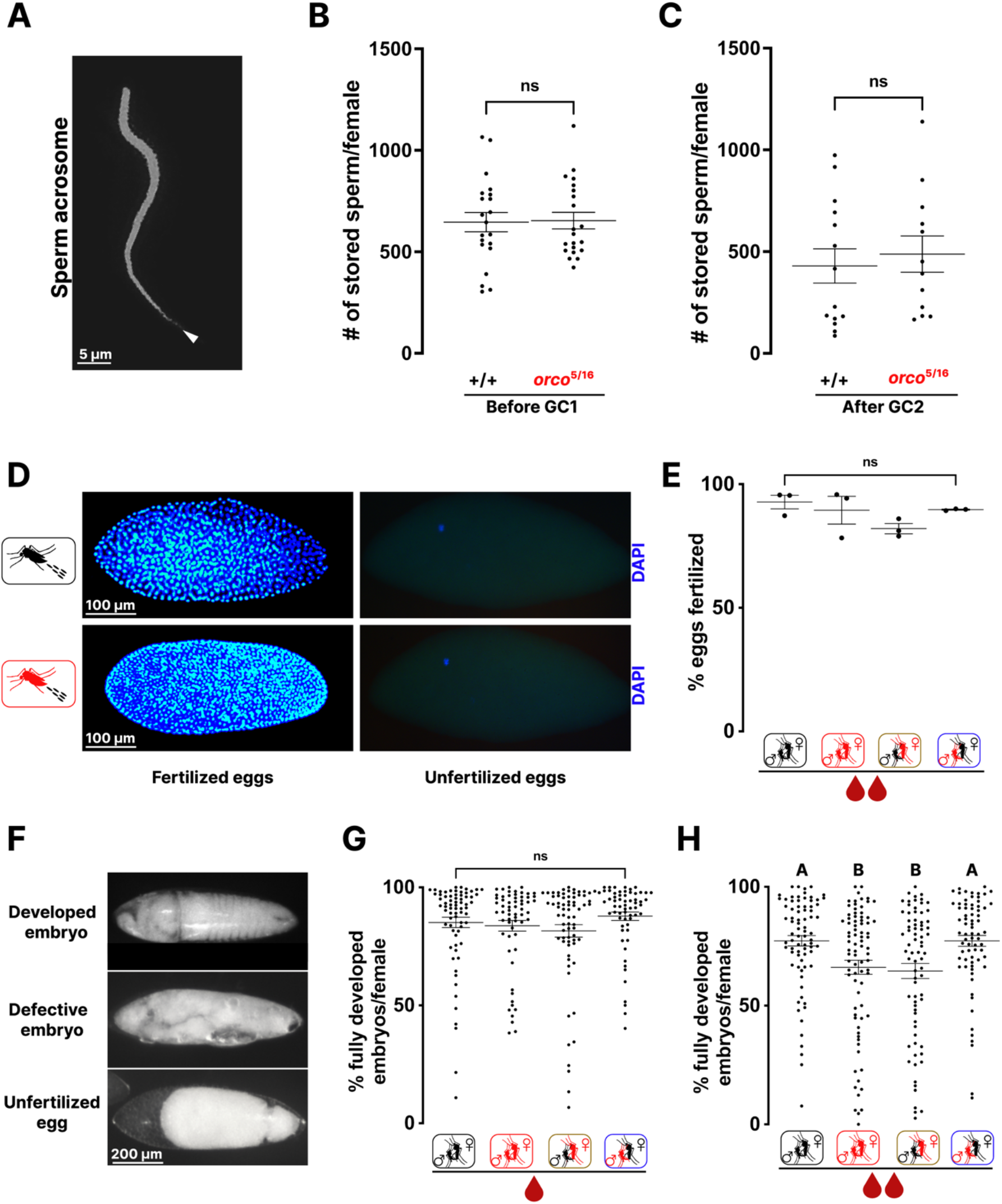
An *orco*-dependent maternal contribution is required for optimum embryonic development over multiple gonotrophic cycles in *Ae. aegypti*. **(A)** Flourescence microscope image of *Ae. aegypti* sperm acrosome stained with DAPI. White arrowhead indicates sperm head. **(B)** Number of sperm stored by individual female wild-type and *orco^5/16^* mosquitoes before going through the first gonotrophic cycle (n=21, p=0.9603), and **(C)** after the second gonotrophic cycle was completed (n=12-14, p=0.5604). **(D)** Flourescence microscope images of DAPI stained fertilized and unfertilized eggs from female mosquitoes of indicated genotypes in the second gonotrophic cycle. **(E)** Percentage of fertilized eggs from individual female mosquitoes of indicated mating groups in the first and second gonotrophic cycles (n=3, p=0.2775). **(F)** Percentage of fully developed embryos produced by individual female mosquitoes of indicated mating groups in the first gonotrophic cycle (n=72-81, p=0.3230). **(G)** Representative light microscope images of developed or defective embryo and unfertilized egg cuticle preparations from wild-type and *orco* mutant females in the second gonotrophic cycle. **(H)** Percentage of fully developed embryos produced by individual female mosquitoes of indicated genotypes & mating groups in the second gonotrophic cycle (n=61-70, p=0.0014). For all dot plots, the long line represents the mean and short lines represent standard error. Statistical analysis for B & C was performed using Mann-Whitney test, and for E, F & H using Kruskal-Wallis test with Dunn’s multiple comparisons. Groups marked with different letters are significantly different. ns indicates no significant difference between comparisons.

We next asked if the sperm stored by *orco^5/16^* females retains the ability to fertilize eggs over multiple gonotrophic cycles. *Ae. aegypti* eggs are fertilized within the first hour after being deposited, and a syncytial blastoderm is formed by 6 hours post egg laying ^73^. Within the first hour of laying, both *Ae. aegypti* and *Drosophila* eggs complete the second meiotic division, leaving each egg with two or three polar nuclei, and a female pronucleus capable of being fertilized in the presence of a male pronucleus ^73,74^. To determine the fertilization status of deposited eggs, pre-blastoderm embryos (2-3 hours after egg laying) were prepared as previously described ^75^, and nuclei were stained with DAPI. Indeed, DAPI treatment of unfertilized eggs usually shows a single nucleus (Figure 3D) ^74^. We found no significant difference in the proportion of fertilized eggs in the second gonotrophic cycle across the four mating groups (Figure 3E).

Our egg fertilization results led us to hypothesize that embryos produced during the second gonotrophic cycle that fail to hatch possibly have a developmental defect. To test this hypothesis, we examined the developmental state of embryos at the end of embryogenesis. Prior to hatching, the characteristic feature of a fully developed *Ae. aegypti* embryo is a segmented external morphology with a clearly distinguishable head, thorax, and abdomen at 77.5 to 96 hours post egg laying ^5,73,76,77^. To determine the developmental status of embryos from wild-type and *orco^5/16^* females, we performed embryo cuticle preparations as previously described ^78^. Independent of the genotype of the male with which they were mated, *orco^5/16^* females produced a significantly reduced number of fully developed embryos in the second gonotrophic cycle compared to controls (Figure 3F-H). Additionally, many of these embryos failed to complete development (Figure 3F).

### CRISPR-based repair of the *Ae. aegypti orco* allele alleviates the fertility decline

Given the reduced egg hatching rate in the second gonotrophic cycle of *orco^5/16^* females, we wondered if repairing the mutation in the *orco* mutant allele would be sufficient to revert the fertility phenotype. Using CRISPR/Cas9 gene editing, we repaired the *orco* allele by restoring the deleted 16bp into the *orco^16^* locus (Figures 4A, 4B & S4A) ^79^. We integrated a single-stranded DNA oligo donor containing a synonymous version of the previously deleted endogenous 16 bases flanked by 92 bases of homologous sequence surrounding the CRISPR binding site into the *orco^16^* locus to generate an *orco-revertant* (*orco-rev*) allele. Six synonymous polymorphisms were included in the reconstituted 16 bases to mark the revertant allele (Figure S4A). We designed an sgRNA that specifically targets the *orco^16^* allele, with a predicted CRISPR cut site 6 bp downstream of the deletion site. An injection mix of sgRNA, Cas9 protein and the repair ssODN was delivered into pre-blastoderm embryos (Figure 4B). Full-length donor DNA sequence was detected by Sanger sequencing of PCR products obtained from primers that anneal outside the bounds of the donor sequence. Sanger sequencing of heterozygous *orco-rev^16^* mosquitoes confirmed ssODN integration (Figures S4B & S4D). To reduce the possibility of off-target effects while ensuring that the *orco^16/16^* genetic background is retained, heterozygous *orco-revertant* mosquitoes were outcrossed to *orco^16/16^* for three generations and then homozygosed (Figures S4C & S4E). We found no apparent deficiency in the overall fitness of *orco-rev^16^* mosquitoes that could suggest their unsuitability for experimental analysis.

**Figure 4.**
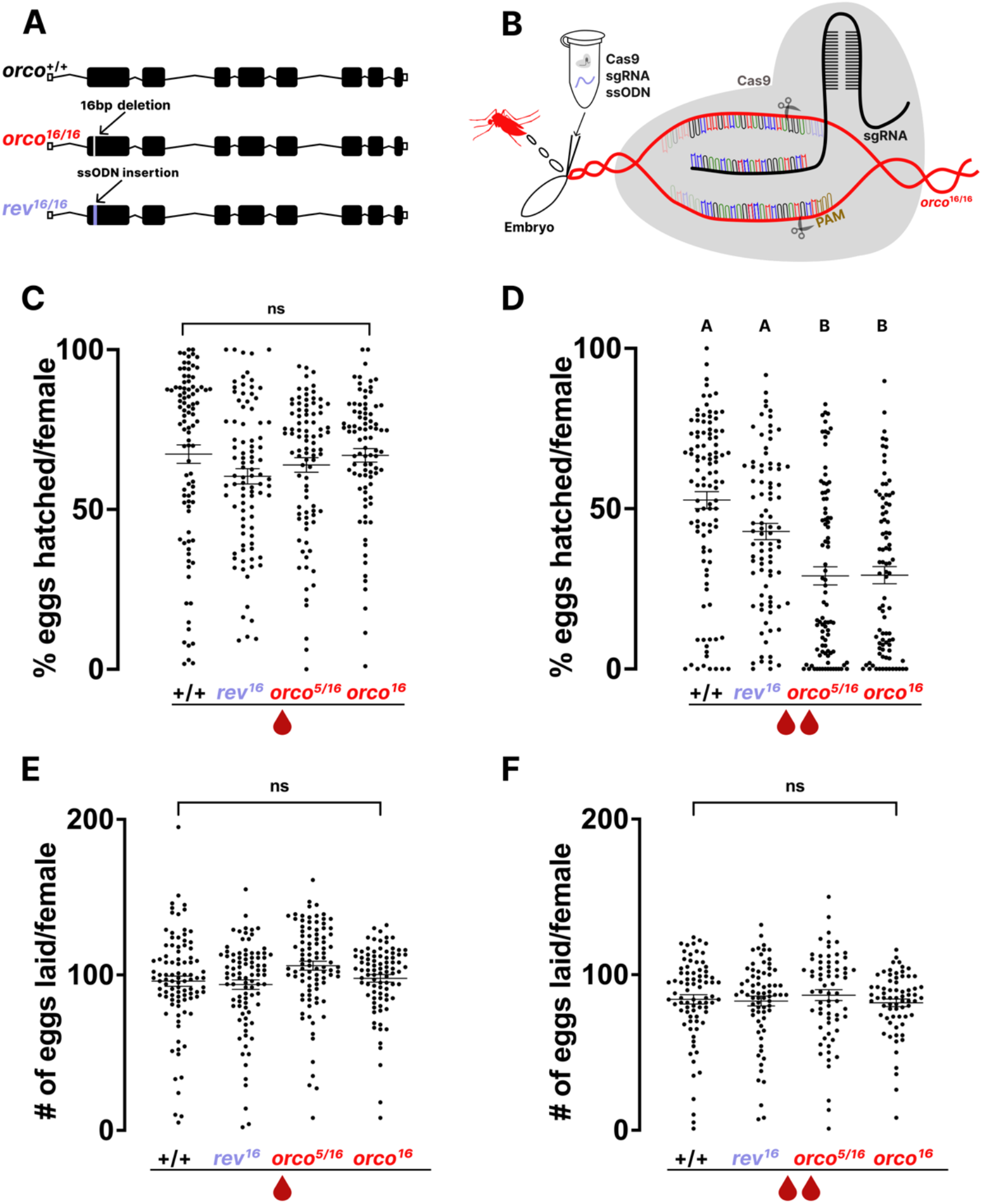
Reconstitution of a wild-type *orco* locus in the *orco* mutant *Ae. aegypti* reverts fertility phenotype. **(A)** CRISPR sgRNA designed to target the Cas9 nuclease of the *orco^16^* mutant locus. Integration location of the ssODN containing the deleted 16bp in exon 1 to repair the mutation. **(B)** Schematic of CRISPR reagents microinjected into pre-blastoderm embryos and Cas9 nuclease activity. Guide RNA and sgRNA targeting sequence are color coded to represent the actual sequences. **(C)** Percentage of eggs hatched from individual female mosquitoes of the indicated genotypes in the first (n=86-102, p=0.1445), and **(D)** second gonotrophic cycles (n=86-93, p<0.0001). **(E)** Number of eggs laid by individual female mosquitoes of indicated genotypes in the first (n=87-102, p=0.053), and **(F)** second gonotrophic cycles (n=86-93, p=0.4189). For all dot plots, the long line represents the mean and short lines represent standard error. Statistical analysis was performed using Kruskal-Wallis test with Dunn’s multiple comparisons. Groups marked with different letters are significantly different. ns indicates no significant difference between comparisons.

We next assessed the reproductive phenotype of *orco-rev^16^* mosquitoes by comparing their fecundity and fertility in parallel to wild-type, *orco^16/16^*, and *orco^5/16^* over two gonotrophic cycles. There was no significant difference in egg hatching rate across all genotypes in the first gonotrophic cycle (Figure 4C). In the second gonotrophic cycle, the reduced egg hatching phenotype of *orco* mutant females was reverted by *orco-rev^16^* mosquitoes (Figure 4D). As expected, we found no significant difference in the number of eggs laid by *orco-rev^16^* females in the first or second gonotrophic cycle compared to wild-type and *orco^16/16^* mutant controls (Figures 4E and 4F).

### RNA-seq analysis of pre-zygotic transition *Ae. aegypti* embryos reveals transcriptional changes cluster by gonotrophic cycle and maternal genotype

Previous transcriptomic studies have reported the expression of *orco* in the ovaries of blood-fed and non-blood-fed *Ae. aegypti* females ^54,80^, as well as during embryonic development ^54^. Given that embryos produced by *orco^5/16^* females in the second gonotrophic cycle carry a maternal effect phenotype leading to reduced hatching, we wondered if the loss of *orco* changes the overall transcriptional signature in pre-zygotic transition embryos. To determine this, we profiled gene expression in these embryos in the first and second gonotrophic cycles across the four mating groups in our previous experiments (Figure 5A). In *Ae. aegypti*, the first zygotic transcription occurs 2 hours post egg laying ^54,81,82^. Given this, we collected embryos at 1-2 hours post egg-laying for our RNA-seq study. We characterized the transcriptomes of the four mating groups in the first and second gonotrophic cycles in triplicate. In total, we sequenced each of the 24 stranded RNA-seq libraries to a depth of at least 200 million, 150 bp paired-end reads.

**Figure 5.**
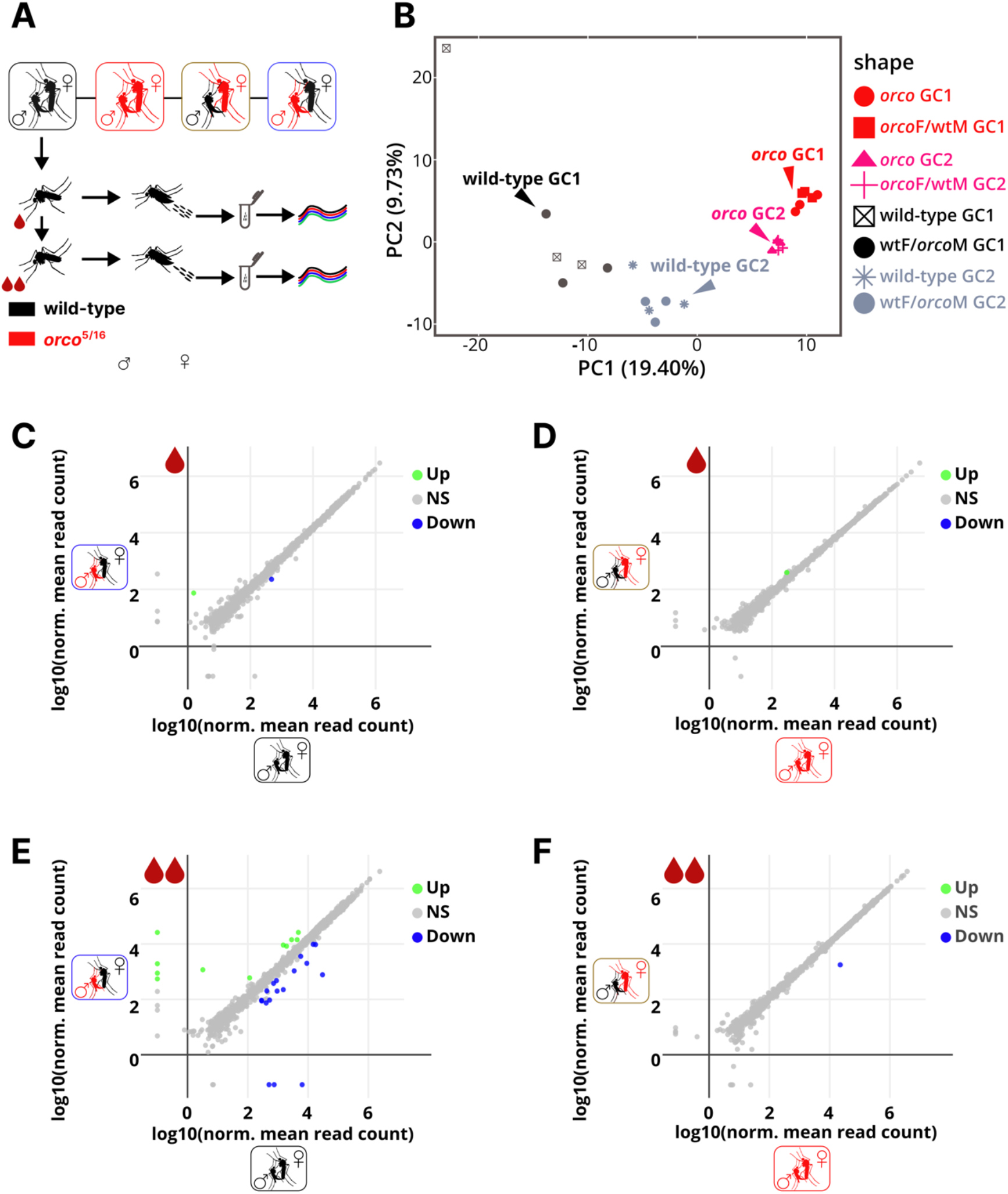
Transcriptional changes cluster by gonotrophic cycle and maternal genotype in the pre-zygotic transition *Ae. aegypti* embryos. **(A)** Schematic of mating groups and sample collection for RNA-seq analysis of 1-2 hours old mosquito embryos in the first and second gonotrophic cycles. **(B)** Principal component analysis of transcriptome-wide expression profile of first and second gonotrophic cycle embryos from the 4 genotypic mating groups. **(C & E)** Differential expression scatter plots comparing log_10_ (Normalized mean) values of read counts in embryos from the two mating groups of wild-type, or **(D & F)** *orco^5/16^* maternal genotypes in the first and second gonotrophic cycles (n=3, FDR <0.01). Statistical analysis for C-F was performed using Wald test. NS represents 10% of genes without significant differential expression. Up and Down represent genes with increased or decreased expression, respectively.

We performed a principal component analysis (PCA) of all libraries to assess clustering patterns among samples ^80^. PCA revealed sample clustering by maternal genotype and gonotrophic cycle across libraries; the libraries from *orco* females were the most tightly clustered (Figure 5B). A similar clustering pattern emerges when normalized reads from all libraries were examined in Euclidean space (Figure S5A). Altogether, these unsupervised hierarchical clustering trends exclude the likelihood of large batch effects or sample cross-contamination during library preparation. Using a weighted gene correlation network analysis (WGCNA) ^83^, we identified gene modules with varied expression levels across treatment groups (Figure S5B). Consistent with our hierarchical cluster results, maternal genotype and gonotrophic cycle clusters are the most represented in our WGCNA modules (Figure S5B). Notably, modules blue and turquoise represent the two most robust clusters and encapsulate genes whose expression levels are regulated by maternal genotype across samples (Figures S5B and S5C).

To confirm that the transcriptional sampling points we assessed are prior to MZT, are dependent on maternal genotype, and are independent of male genetic contribution, we performed differential gene expression analyses between samples from mating groups of the same maternal genotype mated either to wild-type or *orco^5/16^* males, for two gonotrophic cycles (Figure 5A). We found almost no differentially expressed genes in embryos from the same maternal genotype in the first (Figures 5C and 5D) or the second gonotrophic cycle (Figures 5E and 5F). These results clearly validate that the transcriptional signal in our libraries is maternally derived as the genetic contribution of the male had no effect.

### Transmembrane ion channel transcripts are differentially regulated in *orco* mutant *Ae. aegypti* early embryos

We next performed differential expression analyses between wild-type and *orco* mutant female derived libraries. For each pairwise comparison between maternal genotypes, we found over 1,200 differentially expressed genes (DEGs) in the first (Figures 6A, 6C and 6E) and second (Figures 6B, 6D and 6F) gonotrophic cycles. Furthermore, genes differentially expressed in *orco^5/16^* female background contain more than twice as many downregulated genes than upregulated genes across treatment groups (Figures 6A-6F). These unexpectedly high transcriptional changes resulting from mutation in the *orco* gene suggests a regulatory function for *orco* in the female mosquito germline.

**Figure 6.**
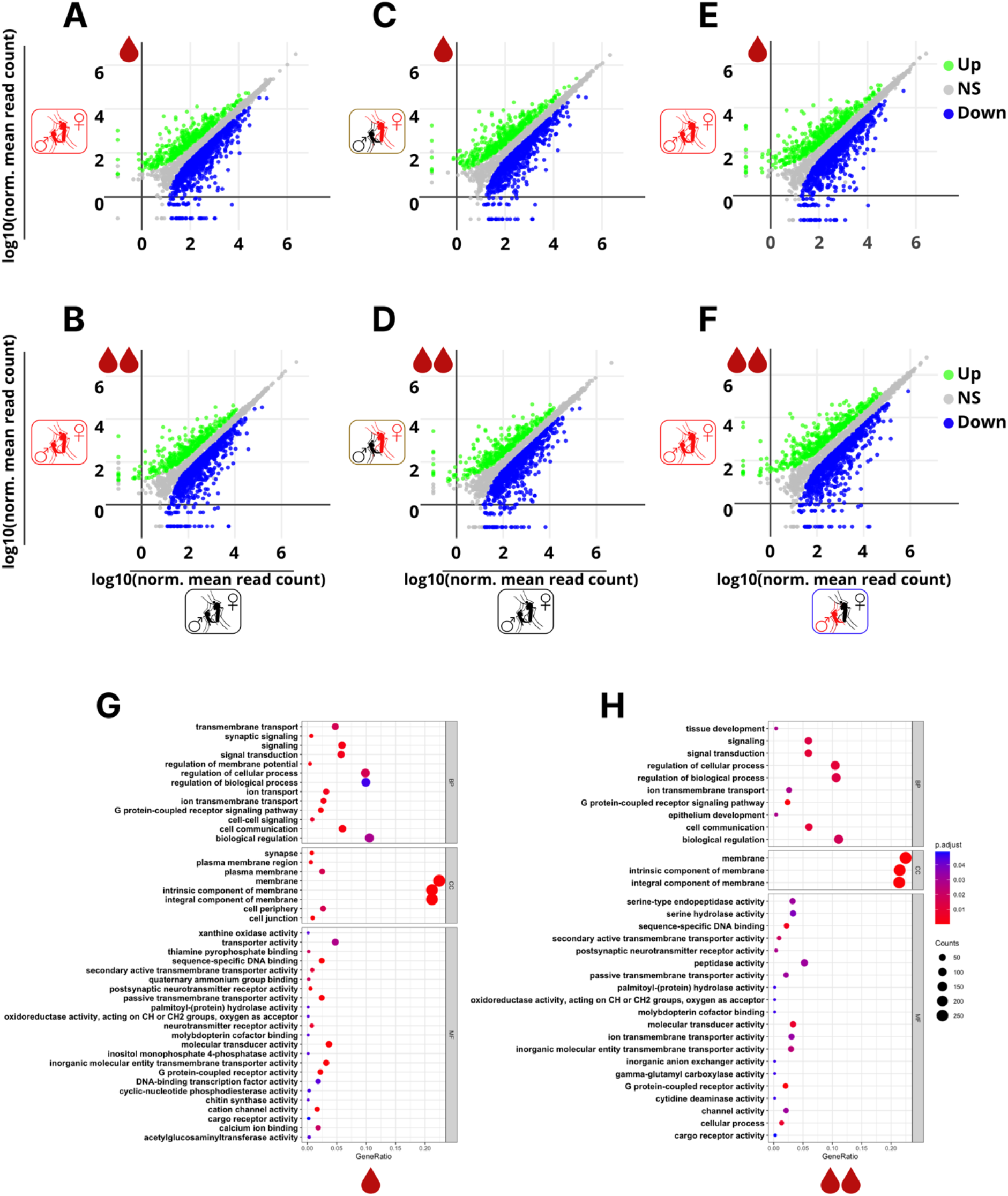
Transmembrane ion channel signaling is robustly represented among differentially expressed genes between wild-type and *orco* mutant *Ae. aegypti* embryos. Differential expression scatter plots comparing log_10_ (Normalized mean) values of read counts in embryos from wild-type and *orco^5/16^* females mated respectively to males of the same genotype **(A & B)**, wild-type females mated to wild-type males & *orco^5/16^* females mated to wild-type males **(C & D)**, and wild-type females mated to *orco^5/16^* males & *orco^5/16^* females mated to *orco^5/16^* males **(E & F)**, in the first and second gonotrophic cycles (n=3, FDR<0.01). **(G)** Enriched gene ontology terms among differentially expressed genes in (A), FDR <0.05. **(H)** Enriched gene ontology terms among differentially expressed genes in (B), FDR 0.05. Statistical analysis for A-F was performed using Wald test. Statistical significance for G & H was determined using Benjamini-Hochberg Procedure. NS represents 10% of genes without significant differential expression. Up and Down represent genes with increased or decreased expression, respectively.

We next assessed the DEGs for enrichment of gene ontology terms across molecular function, biological process, and cellular component classes. Transmembrane ion channel signaling is the most enriched functional trait among the DEGs in embryos from wild-type and *orco^5/16^* females (Figures 6G and 6H). A consistent robust representation of membrane signaling components was obtained in the DEGs for all other cross-maternal genotype pairwise comparisons in both gonotrophic cycles (Figures S6A-D). As the *orco* gene encodes a transmembrane ion channel protein ^53^, these results suggest that *orco* could act in concert with other ion channels during late oogenesis or early embryogenesis to ensure mosquito eggs develop and hatch.

## DISCUSSION

Here, we present a novel non-olfactory phenotype for an olfactory co-receptor in *Ae. aegypti* female mosquitoes. Our evidence strongly suggests that *orco* is required by *Ae. aegypti* females to sustain fertility over multiple gonotrophic cycles. We demonstrate that *orco* mutant females are as capable of ingesting blood meals, storing sperm, and producing fertilized eggs as wild-type females across multiple gonotrophic cycles. Eggs produced by *orco* mutant females after the first gonotrophic cycle show significantly reduced hatching despite no detectable loss in fecundity when compared to wild-type females. We propose that this is likely due to a pre-zygotic requirement for *orco* that promotes embryonic development and hatching. Results from our experiments with revertant mosquitoes strongly suggest that the loss of *orco*, not off-target mutations in the *orco* mutant background, causes this fertility phenotype.

Our hatching assays with cross-genotype mated females strongly suggest that *orco* is a maternal effect gene which acts through the germline to ensure sustained fertility, with no apparent contribution from the male. Our findings argue against the hypothesis that *orco* acts through the male germline to enhance reproductive success by facilitating sperm orientation leading to fertilization in mosquitoes ^55^. *orco* may affect sperm behavior, but it is not altering their storage in the female or their ability to fertilize eggs in our assays.

Oocyte activation is an important prerequisite that triggers a cascade of events, including the resumption of meiosis, translation of maternally transcribed genes, and protein modification or degradation, ultimately preparing the egg for embryonic development ^84^. The flow of Ca^2+^ into intracellular spaces is a critical component of an egg activation event. Although Ca^2+^ wave initiation is inhibited in *Drosophila trpm* null mutants, this inhibition does not prevent egg activation from taking place ^44^. This suggests that egg activation may be under the control of multiple maternal genes. Given that *orco* encodes a membrane ion channel, the defect seen in embryos from *orco* mutant females may be due to reduced calcium flux that does not prevent the egg activation necessary for fertilization. Alternatively, Orco function may be required during early embryogenesis.

Calcium ion channel activity is involved in embryo development ^85,86^. The *Drosophila* KCNQ (*dKCNQ*) is a calmodulin binding voltage-dependent channel expressed in the nurse cells of ovaries, oocytes, pre-zygotic transition embryos, as well as in the fly brain ^87^. Embryos produced by homozygous *dKCNQ* mutant females showed disorganized mid-cleavage nuclei, and consequently, failed to hatch. 44 candidate ion channel genes have been identified to function in *Drosophila* morphogenesis ^88^, several of which belong to the bestrophin, pickpocket (*ppk*), and ionotropic receptor (IR) gene families, including one odorant receptor (OR47a). Analysis of loss-of-function and RNAi transgenic lines for these genes reveal developmental phenotypes impacting the wing, but their role in embryogenesis has not been explored ^88^. In addition, IP_3_-mediated calcium signals are released from the ER during mitotic divisions in syncytial *Drosophila* embryos and regulate the nuclear cycle in early development ^89^. Consequently, a knockout of the IP_3_ receptor disrupts embryonic and larval development ^89^. Together, this evidence suggests that contributions from multiple ion channels are likely required during embryogenesis.

The nearly exclusive representation of transmembrane ion channel processes in our transcriptomic analysis supports the hypothesis that *orco* is a regulator of ion channel signaling in *Ae. aegypti* embryos. Orco could be a key player in a cascade of calcium-dependent signaling events that promote embryogenesis. Further study will be necessary to connect *orco* to other molecular targets that promote the development and hatching of mosquito embryos. Finding connections between these molecular targets may yield new approaches for mosquito population control.

## Supporting information

Sequencing Counts

## ACKNOWLEDGEMENTS

We are grateful for the comments and support from the DeGennaro laboratory on the manuscript. We thank Fredis Mappin for his help by sharing scripts to help our bioinformatic analysis. We thank Alejandro Acuna for his comments on the manuscript. O.G.D. was supported by the Florida International University Biological Sciences Department. This research was supported in part by the United States Centers for Disease Control (CDC) Grant 1U01CK000510: Southeastern Regional Center of Excellence in Vector-Borne Diseases: The Gateway Program. The CDC did not have a role in the design of the study, the collection, analysis, or interpretation of data, nor in writing the manuscript.

## AUTHOR CONTRIBUTIONS

O.G.D. and M.D. designed the research. O.G.D., K.M.S., A.V.A., and A.L.C.S. performed the experiments. O.G.D., A.L.C.S., and A.J.B. analyzed the data. O.G.D. and M.D. wrote the paper.

## SUPPLEMENTARY FIGURES

**Figure S1.**
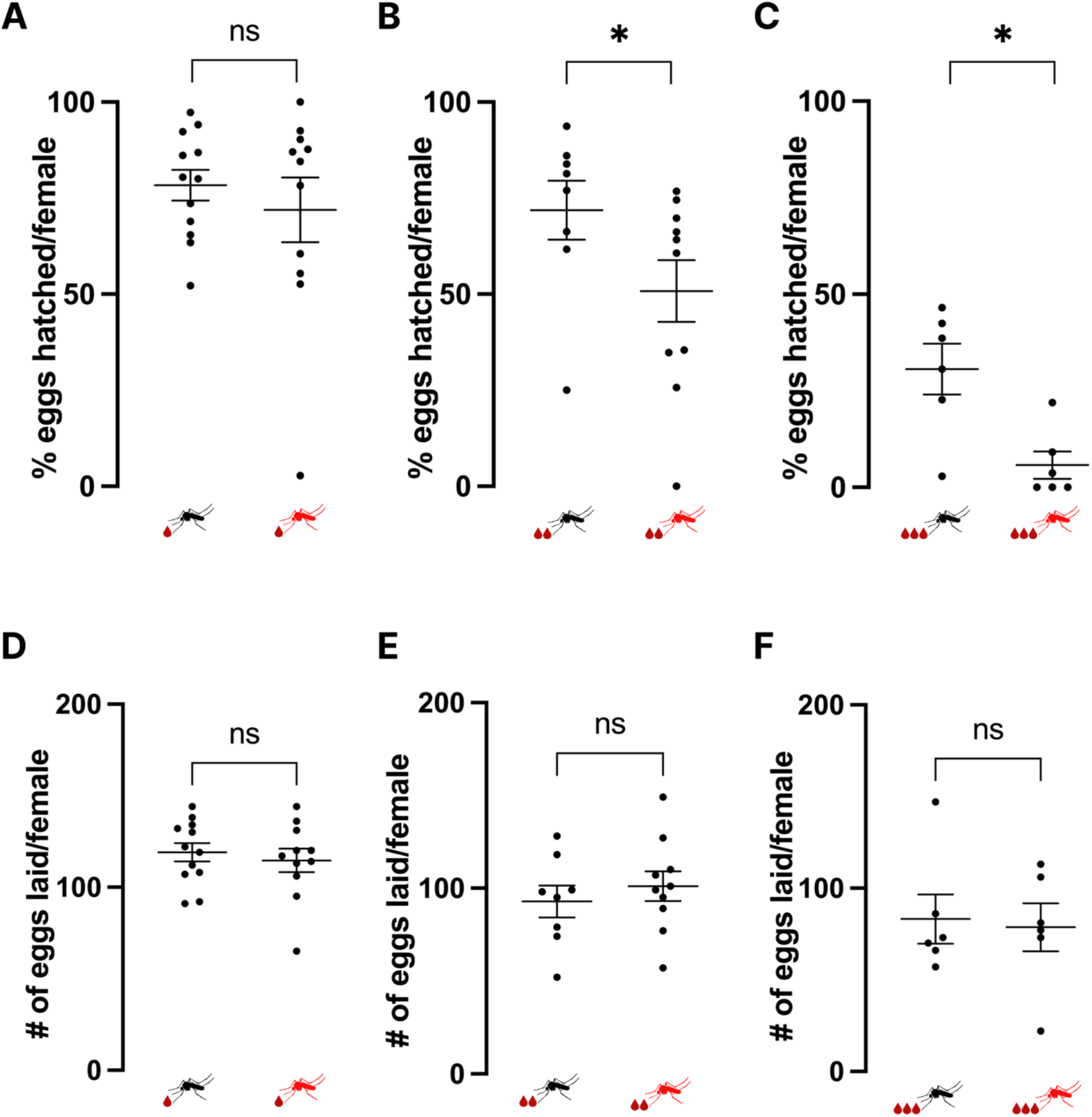
Fertility and fecundity of wild-type and *orco* mutant *Ae. aegypti* female mosquitoes across three gonotrophic cycles. **(A)** Percentage of individual wild-type and *orco^5/16^* female mosquitoes eggs hatched in the first (n=11-12, p=0.4868), **(B)** second (n=8-10, p=0.0434), and **(C)** third (n=6, p= 0.0152) gonotrophic cycles. **(D)** Number of eggs laid by individual wild-type and *orco^5/16^* female mosquitoes in the first (n=11-12, p=0.5889), **(E)** second (n=8-10, p= 0.4967), and **(F)** third (n=6, p=0.8149) gonotrophic cycles. A-C was analyzed using Mann-Whitney test, and D-F using unpaired t-test. Groups marked with asterisks (*) are significantly different (p < 0.05). ns indicates not significant.

**Figure S2.**
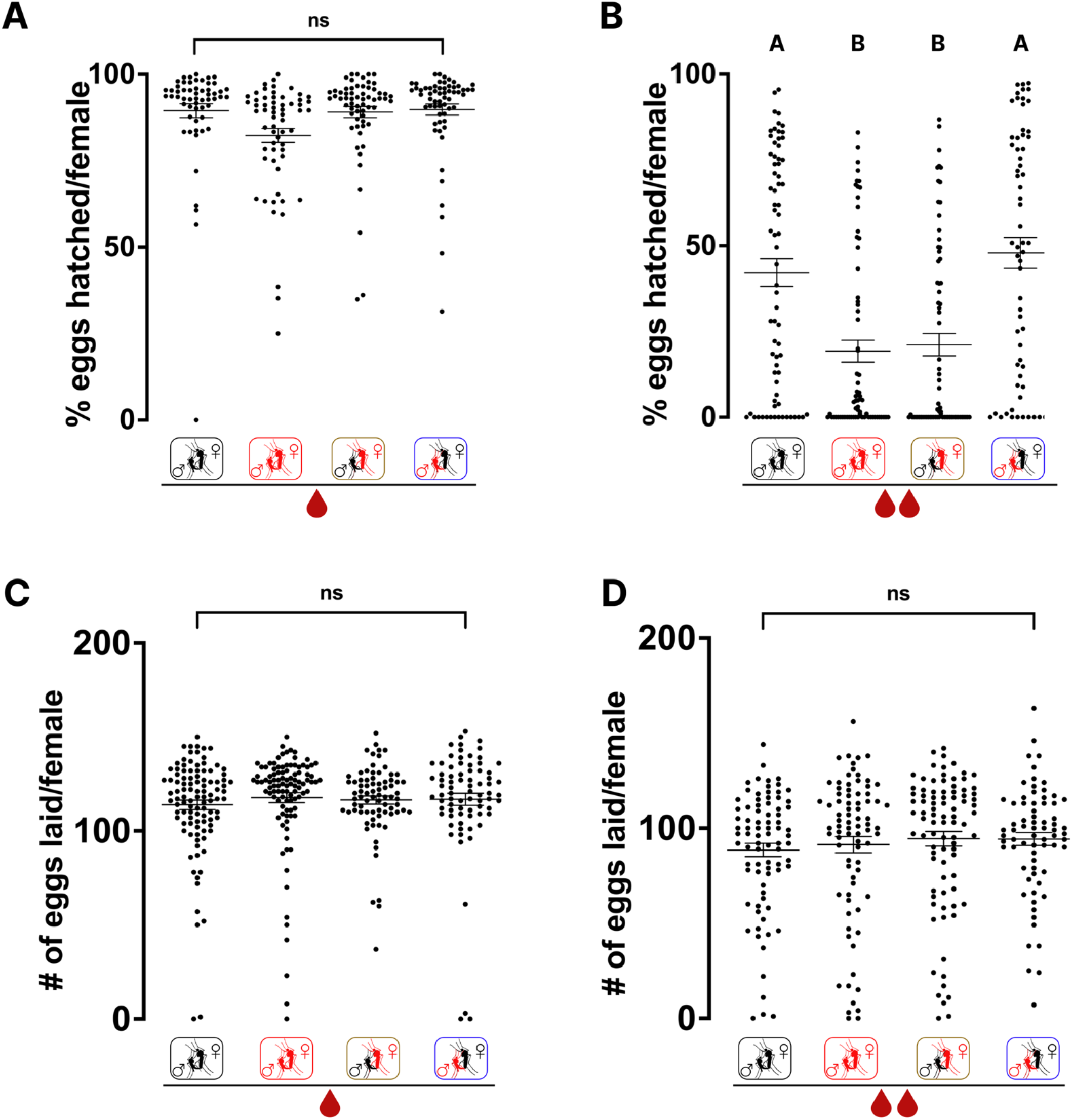
A second gonotrophic cycle maternal effect fertility phenotype in *orco* mutant *Ae. aegypti* mated in pools. **(A)** Percentage of individual female mosquitoes of indicated genotypes & mating groups eggs hatched in the first (n=63-72, p=0.0606), and **(B)** second gonotrophic cycles (n=57-62, p<0.0001). **(C)** Number of eggs laid by individual female mosquitoes of indicated genotypes & mating groups in the first (n=74-77, p=0.0752), and **(D)** second gonotrophic cycles (n=63-72, p=0.2509). Mosquitoes were allowed to freely mate in a cage. For all dot plots, the long line represents the mean and short lines represent standard error. All the data above were analyzed using Kruskal-Wallis test with Dunn’s multiple comparisons. Groups marked with different letters are significantly different. ns indicates no significant difference between comparisons.

**Figure S4.**
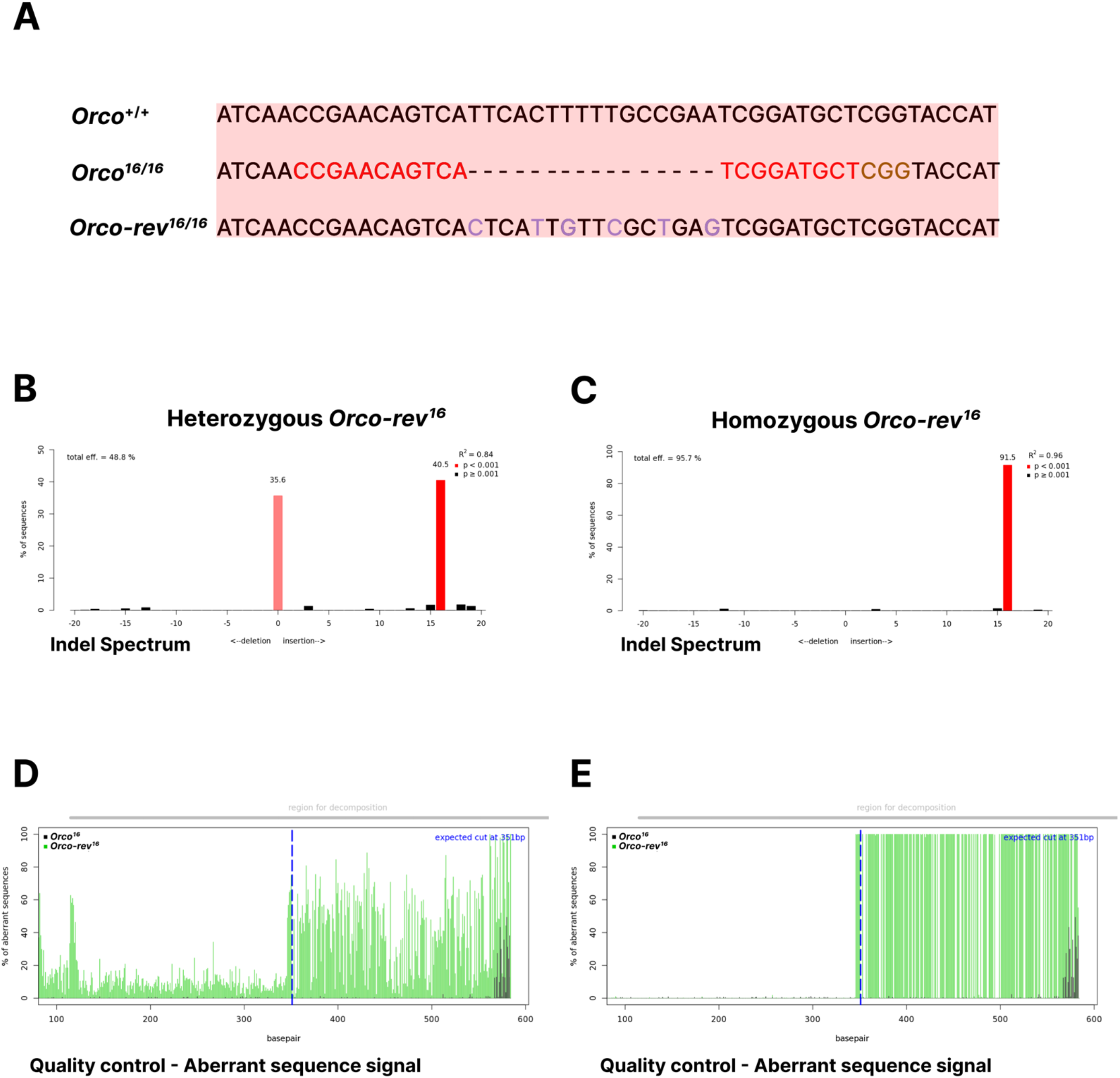
Validation of ssODN insertion using TIDE analysis. **(A)** *orco* deletion site showing an integrated 16bp in the *orco-rev^16^* allele. sgRNA targeting sequence (red), PAM sequence (gold) and SNPs (purple) **(B)** *orco^16^* and *orco-rev^16^* alleles in heterozygous *orco-rev^16^* mosquito. **(C)** *orco-rev^16^* allele in homozygous *orco-rev^16^* mosquito. **(D)** *orco^16^* and *orco-rev^16^* sequence signals around the CRISPR site in heterozygous, and **(E)** homozygous mosquitoes, respectively. Significance cutoff for decomposition at p=0.001.

**Figure S5.**
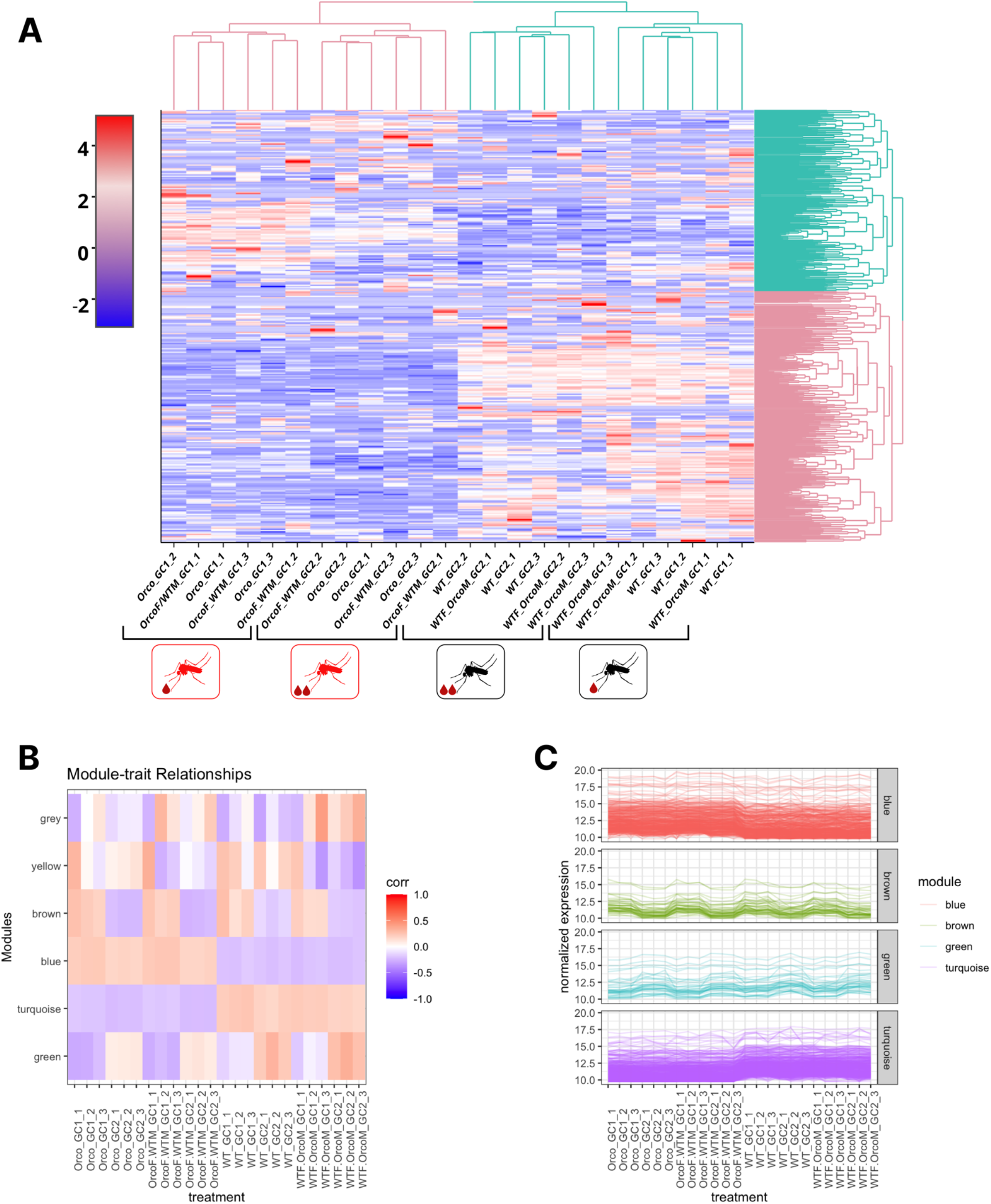
Transcriptome wide gene clusters by gonotrophic cycle and maternal genotype. **(A)** Transcriptome-wide heat map representation of normalized reads showing sample clusters (column dendrograms) by gonotrophic cycle and maternal genotype. **(B)** Weighed gene correlation network analysis (WGCNA) showing gene modules (clusters) by reads signal levels across samples. **(C)** Signal levels of genes within selected WGCNA modules across samples.

**Figure S6.**
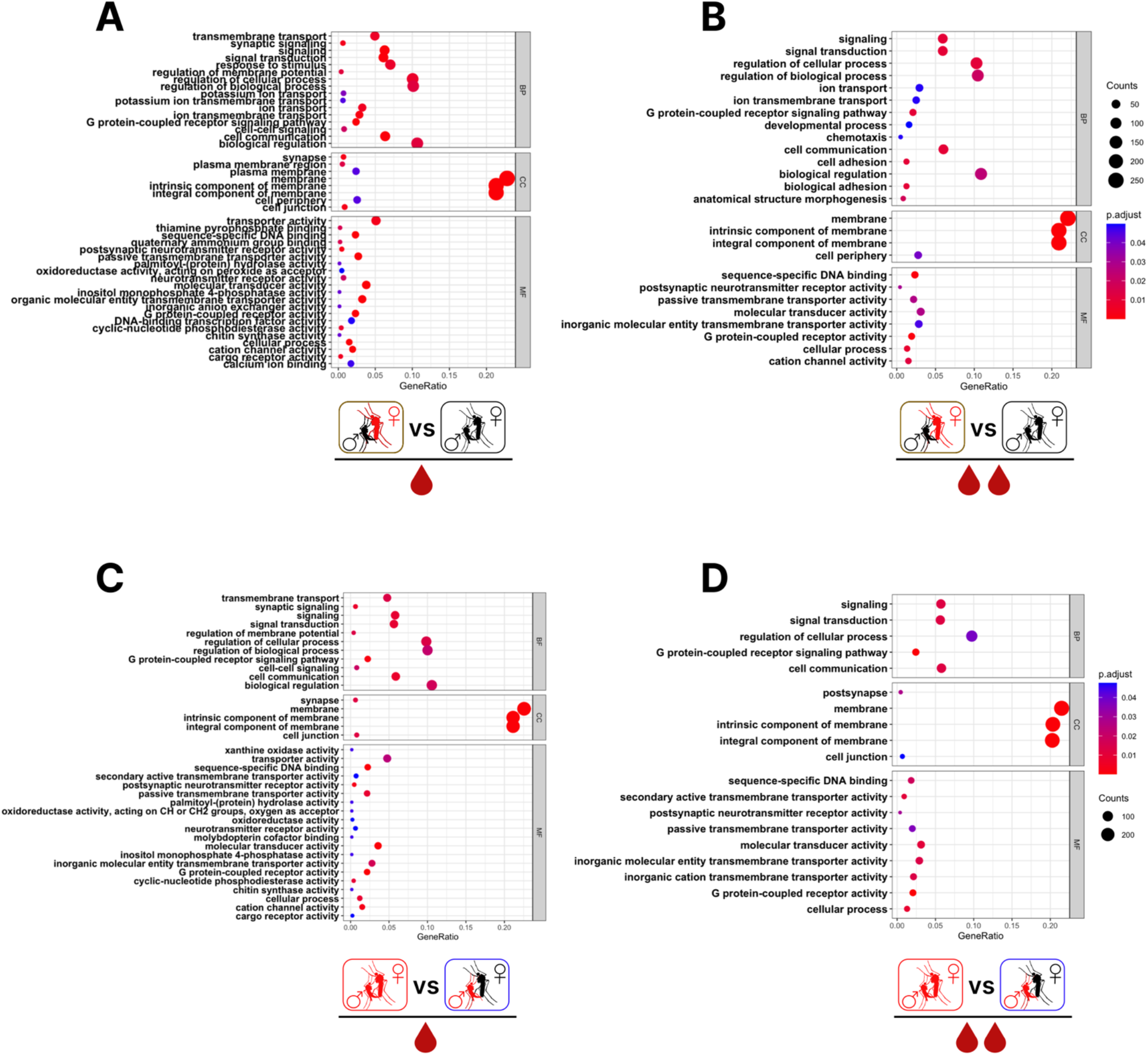
Gene ontology enrichment among differentially expressed genes. **(A & B)** Enriched gene ontology among DEGs in Fig. 6A & 6B, respectively. **(C & D)** Enriched gene ontology among DEGs in Fig. 6C & 6D, respectively. Statistical significance was determined using Benjamini-Hochberg Procedure, FDR<0.05.

## KEY RESOURCES TABLE

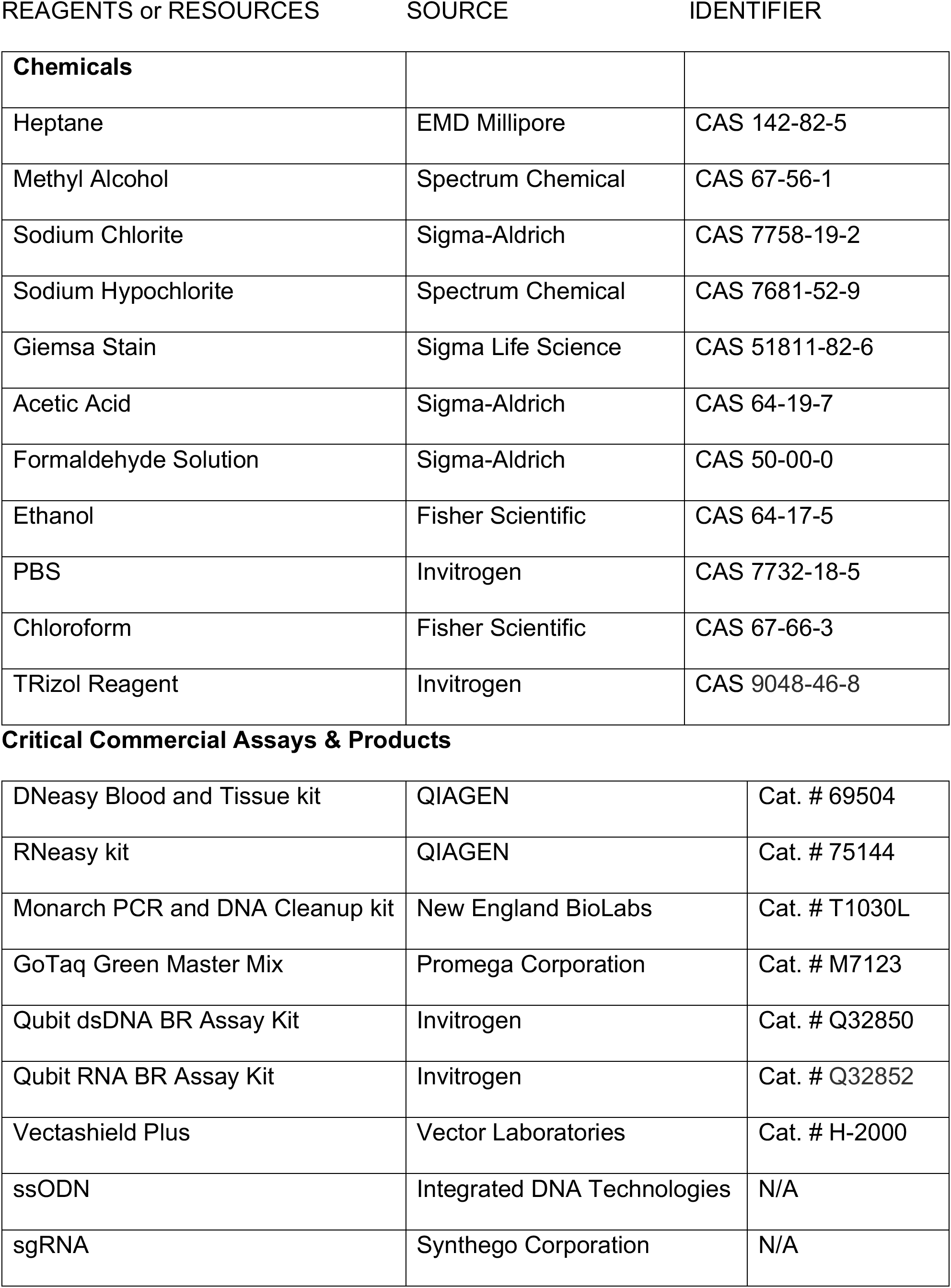
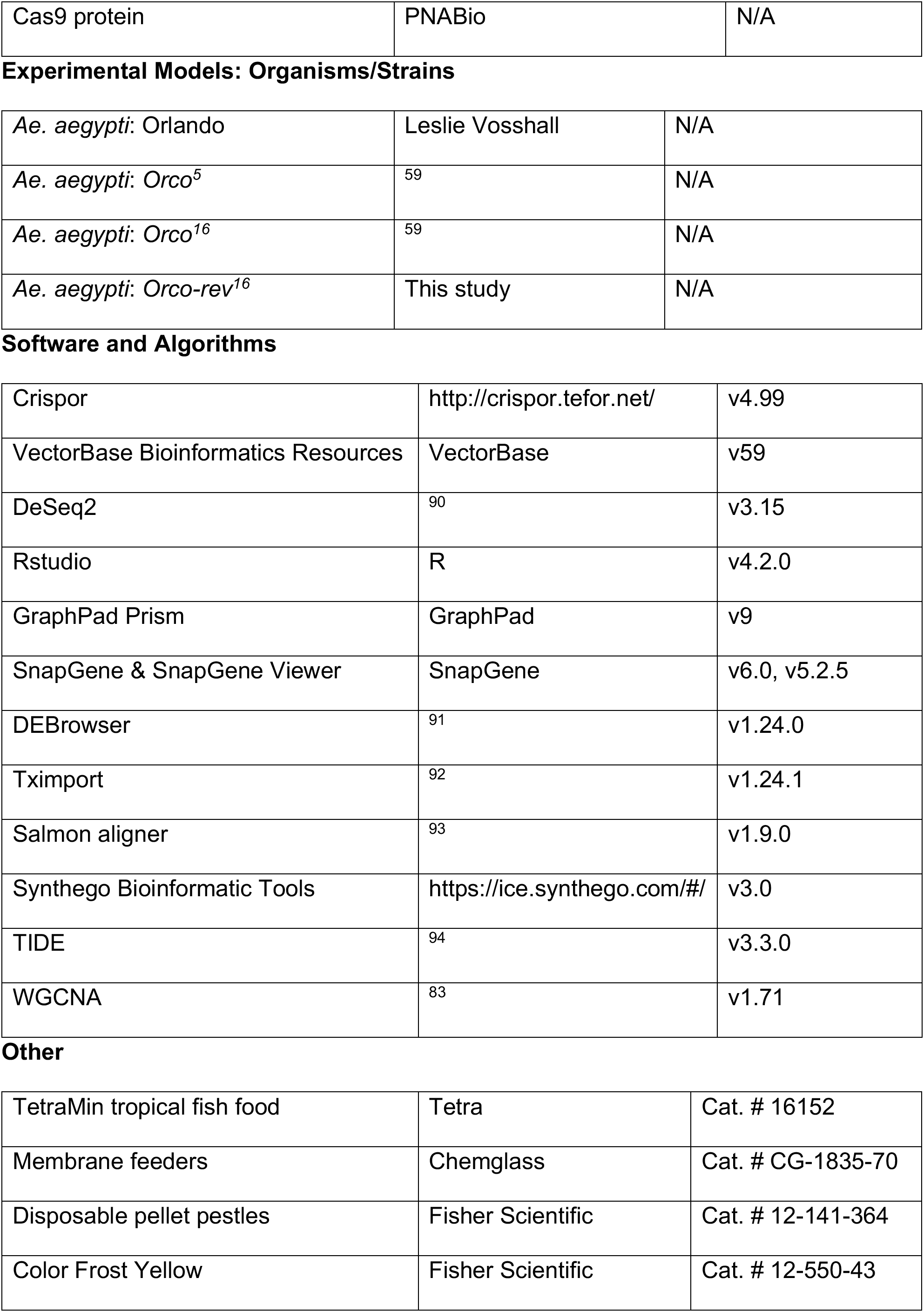
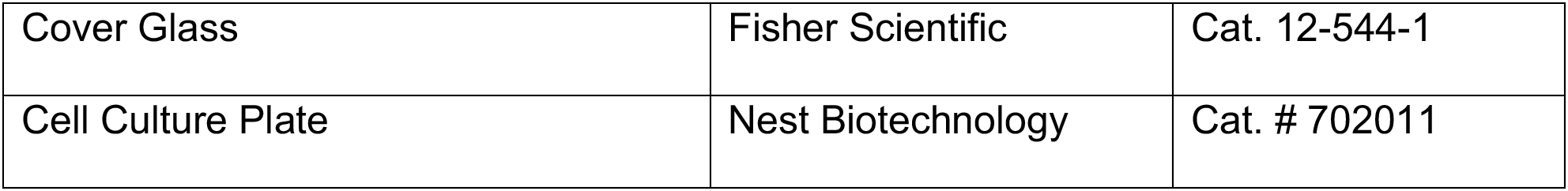

IDENTIFIER
| **Chemicals** |  |  |
| --- | --- | --- |
| Heptane | EMD Millipore | CAS 142-82-5 |
| Methyl Alcohol | Spectrum Chemical | CAS 67-56-1 |
| Sodium Chlorite | Sigma-Aldrich | CAS 7758-19-2 |
| Sodium Hypochlorite | Spectrum Chemical | CAS 7681-52-9 |
| Giemsa Stain | Sigma Life Science | CAS 51811-82-6 |
| Acetic Acid | Sigma-Aldrich | CAS 64-19-7 |
| Formaldehyde Solution | Sigma-Aldrich | CAS 50-00-0 |
| Ethanol | Fisher Scientific | CAS 64-17-5 |
| PBS | Invitrogen | CAS 7732-18-5 |
| Chloroform | Fisher Scientific | CAS 67-66-3 |
| TRizol Reagent | Invitrogen | CAS 9048-46-8 |
| Critical Commercial Assays & Products |  |  |
| DNeasy Blood and Tissue kit | QIAGEN | Cat. # 69504 |
| RNeasy kit | QIAGEN | Cat. # 75144 |
| Monarch PCR and DNA Cleanup kit | New England BioLabs | Cat. # T1030L |
| GoTaq Green Master Mix | Promega Corporation | Cat. # M7123 |
| Qubit dsDNA BR Assay Kit | Invitrogen | Cat. # Q32850 |
| Qubit RNA BR Assay Kit | Invitrogen | Cat. # Q32852 |
| Vectashield Plus | Vector Laboratories | Cat. # H-2000 |
| ssODN | Integrated DNA Technologies | N/A |
| sgRNA | Synthego Corporation | N/A |
| Cas9 protein | PNABio | N/A |
| Experimental Models: Organisms/Strains |  |  |
| *Ae. aegypti*: Orlando | Leslie Vosshall | N/A |
| *Ae. aegypti*: Orco<sup>5</sup> | 59 | N/A |
| *Ae. aegypti*: Orco<sup>16</sup> | 59 | N/A |
| *Ae. aegypti*: Orco-rev<sup>16</sup> | This study | N/A |
| Software and Algorithms |  |  |
| Crispor | http://crispor.tefor.net/ | v4.99 |
| VectorBase Bioinformatics Resources | VectorBase | v59 |
| DeSeq2 | 90 | v3.15 |
| Rstudio | R | v4.2.0 |
| GraphPad Prism | GraphPad | v9 |
| SnapGene & SnapGene Viewer | SnapGene | v6.0, v5.2.5 |
| DEBrowser | 91 | v1.24.0 |
| Tximport | 92 | v1.24.1 |
| Salmon aligner | 93 | v1.9.0 |
| Synthego Bioinformatic Tools | https://ice.synthego.com/#/ | v3.0 |
| TIDE | 94 | v3.3.0 |
| WGCNA | 83 | v1.71 |
| Other |  |  |
| TetraMin tropical fish food | Tetra | Cat. # 16152 |
| Membrane feeders | Chemglass | Cat. # CG-1835-70 |
| Disposable pellet pestles | Fisher Scientific | Cat. # 12-141-364 |
| Color Frost Yellow | Fisher Scientific | Cat. # 12-550-43 |
| Cover Glass | Fisher Scientific | Cat. 12-544-1 |
| Cell Culture Plate | Nest Biotechnology | Cat. # 702011 |

## EXPERIMENTAL MODEL DETAILS

### Statement of Research Ethics

All research was conducted in compliance with the NIH guidelines and the Florida International University Environmental Health and Safety guideline. All maintenance and experiments with genetically modified strains of *Ae. aegypti* mosquitoes were performed in Arthropod Containment Level 2 (ACL2) facilities. Biohazard disposal, laboratory practices, facilities and equipment were reviewed and approved by the Florida International University Institutional Biosafety Committee (IBC-21-022-AM03)

### Mosquito rearing

Wild-type and mutant *Ae. aegypti* mosquitoes were reared and maintained in the insectary at 27±1°C and 70±10% relative humidity in a 14:10 light-dark cycle with lights on at 8 AM Mosquito eggs were hatched in a vacuum sealed Mason jar containing 0.5 L of pre-boiled deionized deoxygenated water with a dissolved TetraMin tropical fish food tablet (Tetra, Melle, Germany). 250 hatched L2 larvae were sorted into 2 L of deionized (DI) water in a rearing pan (5.3 L polycarbonate food pan, Carlisle, Oklahoma, USA), and fed TetraMin tablets until pupation. Pupae were transferred to a porcelain ramekin containing DI water and placed in a 30 × 30 × 30 cm BugDorm-1 insect rearing cage (MegaView Science Co., Ltd., Taiwan). Emerged adults were maintained in a 1:1 male:female ratio with access to 10% sucrose solution *ad libitum*. To generate eggs for colony maintenance, 5-7-day-old females were blood-fed using a 50mm glass feeder (#1588-50, NDS Technologies, Vineland, NJ) covered with stretched Parafilm M wrapping film (#13-374-10, Bemis, Thermo Fisher Scientific^™^, Waltham, MA) pre-rubbed on human arm to enhance female’s attraction. The feeder was filled with 3 ml of defibrinated whole sheep blood (#R54020, Remel Inc, Thermo Fisher Scientific^™^, Lenexa, KS) pre-warmed to 37°C, and supplemented with 0.4 mM ATP (#34369-07-8, ACROS Organics, Thermo Fisher Scientific, Pittsburgh, PA). The feeder was connected to a circulating water bath maintained at 37°C and females were allowed to access the feeder through the mesh panel of the cage for 1 hour. White porcelain ramekin lined with 55mm diameter Whatman 1 filter paper (#1001-055, Cytiva), and half-filled with DI water was placed into the cage 3 days after the blood meal for the females to lay eggs.

### Mosquito weight determination

Five 5-7-days-old females were randomly chosen from a cage of 250 mosquitoes starved on water overnight, cold anaesthetized and individually weighed on a balance scale (Ahaus Adventurer). These weighed females were thereafter sacrificed while the those remaining in the cage were blood-fed. Following blood feeding, 50 engorged females were sorted into a fresh cage and 5 randomly chosen individuals were cold anaesthetized, individually weighed and sacrificed thereafter. The remaining engorged females were allowed to develop their eggs after which they were prepared for the first gonotrophic cycle egg collection. At the end of the first gonotrophic cycle, these females were starved on water overnight and 5 randomly chosen non-blood-fed and blood-fed females were individually weighted before and after blood feeding, respectively.

### Fecundity and fertility assays

To determine the number of eggs produced by female mosquitoes, females were starved on water overnight prior to being provided with a blood meal as described above. 50 fully engorged females were sorted into a new rearing cage with unlimited access to 10% sucrose solution. 72-96 hours post blood meal, females were cold anaesthetized on ice and prepared for individual oviposition as previously described with slight modifications ^95^. 15 mm Whatman 1 filter paper (#1001-0155, Cytiva) was fitted in each well in a 24-well oviposition plate (oviplate) and 80 ml of DI water was added to serve as an oviposition substrate. Females were individually placed into each well of the oviplate with a pair of tweezers and allowed to recover. Egg collection lasted for 24 hours after, and females that did not lay eggs during this period were removed from the assay. Following the first gonotrophic cycle egg collection, females were released into a new rearing cage, starved on water overnight to allow a second gonotrophic cycle egg collection. Oviposition substrates remained moist for at least 72 hours and were subsequently airdried for additional 3-4 days. Fecundity was determined by counting the number of eggs each female deposited. To determine the fertility of each female, eggs from individual female were hatched 6-7 days after they were laid. Each egg paper was submerged in 50ml of hatching broth in a disposable hatching cup. Eggs were allowed to hatch over a period of 3 days after which larvae were counted to determine hatching rates. Scoring was conducted by an observer who was blinded to the genotype of the eggs.

### Spermathecal sperm count

3–5-day-old females were cold anaesthetized and spermathecae preparation and sperm count was carried out as described in ^96^ with modifications. Two pairs of fine tweezers were used to dissect spermathecae from each female on a slide (Color Frost Yellow, Fisher Scientific) with 50 μl of 1% phosphate buffered solution (PBS) under a dissecting microscope. Intact spermathecae from a female were transferred to 20 μl of PBS on a manually ruled slide, torn open using insect pins to release sperm and vigorously agitated to minimize sperm clumping. Insect pins used for each female’s spermathecal dissection were rinsed with 50 μl of PBS twice on the slide, after which the slide was dried at 65°C on a hot plate. Upon drying, sperm was fixed on slide with 70% ethanol at 65°C for 20 mins. Vectashield Plus Antifade Mounting Medium with DAPI (Vector Laboratories) was applied to sperm on the slide for an hour. Sperm heads were viewed at 400x magnification under a flourescence microscope (DM5500B, Leica Microsystems) and counted by slowly walking across the ruled slide. Scoring was conducted by an observer who was blinded to the genotype of the sperm.

### Embryo cuticle preparation

Eggs were collected from individual gravid female over a period of 24 hours and egg papers kept moist for additional 96 hours to allow for embryogenesis to be completed. Embryo cuticle preparation was done as described in ^78^. Individual egg paper was transferred to a 50 ml plastic container containing 20 ml of the following bleach formulations – 8 g sodium chlorite, NaClO2; 4 ml acetic acid; 1000 ml DI water, for 12 hours. Bleached embryos were imaged and counted under a light microscope (MSV269, Leica Microsystems). Prescence of a discernable head, thorax and abdomen was used as scoring criteria for fully developed embryos ^5,77^. Scoring was conducted by an observer who was blinded to the genotype of the embryos.

### Egg fertilization assay

Eggs were collected from a pool of 50 gravid females from each mating group over a period of 1 hour in the first and second gonotrophic cycles and allowed to embryonate for additional 2 hours. Embryo dechorionation, fixation and endochorion disruption was done as described in ^74^. Following endochorion removal, 100 embryos from each mating group were transferred to a slide and stained with Vectashield Plus Antifade Mounting Medium with DAPI (Vector Laboratories) for 1 hour. Slides were mounted and viewed at 100x magnification under a fluorescence microscope (DM5500B, Leica Microsystems). Eggs with a maximum of 4 nuclei were scored as unfertilized while those with 5 or more nuclei were scored as fertilized in our assay. Scoring was conducted by an observer who was blinded to the genotype of the embryos.

### RNA extraction and sequencing

Eggs were collected from a pool of 50 gravid females from each mating group over a period of 1 hour in the first and second gonotrophic cycles. 800-1000 eggs from each mating group aged at 1-2 hours old were transferred to a 2 ml Eppendorf tube (Eppendorf Biopur) and immediately stored at −80°C until RNA extraction. Samples were manually homogenized using RNase-free pestles (Fisher Scientific) in TRIzol Reagent (Invitrogen) as per manufacturer instructions, through the chloroform extraction step, at which point the aqueous phase was recovered and mixed 1:1 with absolute ethanol. This mixture was then processed using the RNeasy Blood and Tissue Kit (Qiagen) with on-column DNase treatment, as per manufacturer instructions with the omission of lysis in Buffer RLT. Sample concentration was determined using Qubit 4 Fluorometer (Invitrogen). Thereafter, RNA quantity and quality were evaluated using an Agilent BioAnalyzer 2100 and RNA 6000 Pico Kit (Agilent Technologies). Stranded library construction and Illumina sequencing of targeting 200 million paired-end 150 base reads per sample was performed at the University of Miami Center for Genome Technology.

### Expression data processing and analysis

Reads from each library were trimmed to remove adapter sequence ^97^, and thereafter mapped to the AaegL5.0 genome gene models using Salmon v1.8.0 ^93^. Mapping resulted in a range of 194-293 million read pairs mapped per library (Supplement S1). Transcript-level abundance estimates for mapped reads were determined using tximport v1.24 ^92^. Differential expression and principal component analyses were performed on read counts using Deseq2 in R’s shiny DE Browser ^91^. Differentially expressed genes for all pair-wise comparisons were determined by an FDR-corrected cutoff of α < 0.01. Heat map representation of raw count clusters across libraries was generated with DE Browser using the following parameters: dist method (‘Euclidean’), normalization method (‘MRN’) and clustering linkage (‘complete’). Gene co-expression network modules were generated across all libraries using the WGCNA package in R ^83^, following Variance Stabilizing Transformation of raw counts with Deseq2 ^90^. Gene ontology enrichment analysis was performed for all differentially regulated genes per pair-wise comparison using resources on vectorbase.org at an FDR of α < 0.01. Obsolete and redundant terms were manually removed, and data visualization was done in R ^98^.

### CRISPR reagents and target site design

CRISPR single guide RNA (sgRNA) and single-stranded DNA oligodeoxynucleotide (ssODN) were designed following the standard protocols ^79^. The sgRNA sequence (CCGAACAGTCATCGGATGCT) for the repair of *orco^16/16^* allele was chosen by the presence of protospacer-spacer adjacent motif (PAM) with the sequence CGG exactly 9bp downstream of the *orco* deletion site. sgRNA sequence was assessed for potential off-target binding sites using CRISPOR. (http://crispor.tefor.net/crispor.py?batchId=Ui0OpgjaKWkcS4POcaqQ). sgRNA synthesis of a 1.5 nmol scale 100-mer was provided by a commercial vendor (Synthego Corporation), with the chemical modification of the 3 terminal bases at each end of the guide to include 2’ O methyl analogs and 3’ phosphorothioate internucleotide linkages to improve RNA stability. sgRNA stock was resuspended in 150 μl of nuclease-free water and concentrations verified using with Qubit RNA Broad Range assay on 2 μl of stock sgRNA solution.

ssODN synthesis was performed by Integrated DNA Technologies using the 20 nmole ultramer service with standard desalting. The ssODN contains the deleted 16bp in the *orco^16/16^* allele flanked by 92bp of homologous sequences to the target site on both sides: TTCACCCATTCCGTGACCAAGTTCATCTACGTCGCCGTCAACTCGGAACATTTCTACCGCACG CTGGGCATCTGGAATCAACCGAACAGTCA**CTCATTGTTCGCTGAG**TCGGATGCTCGGTACCA TTCGATTGCGTTGGCTAAGATGCGAAAACTGCTGGTCATGGTGATGGTGACTACAGTGCTATCCGTCGTCGGTAA (repair sequence in bold, and SNPs in red: T>C, C>T, T>G, T>C, C>T, and A>G). 20 nmol of ssODN was resuspended in 1.2 ml of nuclease free water, yielding ~1000 ng/μl solution as verified by the Qubit DNA Broad Range assay on 1 μl of resuspended ssODN solution. 50 μg of lyophilized Cas9 protein powder (PNA Bio) was resuspended in 50 μl of nuclease free water to obtain a 1000 ng/μl, which was subsequently diluted to the desired working concentration.

### Generation of *orco* revertant allele (*orco-rev^16^*) and detection

To obtain a stable germline knock-in, CRISPR reagents were injected into the posterior end of preblastoderm embryo as previously described ^99^. Pre-blastoderm embryos made up of syncytium nuclei that will become somatic and germline cells, making this a suitable stage for germline transformation by the CRISPR-Cas9 complex. *Ae. aegypti* pre-blastoderm embryo microinjection was carried out at the Invertebrate Transformation facility at Florida International University using the XenoWorks Digital Microinjector system (Sutter Instrument). The microinjection mixes were formulated as follows: sgRNA (40 ng/μl), Cas9 (300 ng/μl), ssODN (200 ng/μl) were diluted into nuclease-free H2O. A total of 361 *orco* mutant embryonated eggs were injected to target exon 1 of the *orco^16/16^* allele.

Embryos were hatched 5 days after injection and reared to the adult stage. 104 out of the injected 361 embryos hatched with approximately even male:female ratio. A total of 45 G_0_ females were allowed to mate freely with *orco^16/16^* males, and females were blood-fed to generate G_1_ progeny. Gravid G_0_ females were individually placed in an oviplate ^94^ to collect G_1_ eggs. After egg laying, G_0_ females were sacrificed to assess CRISPR activities. Mosquito genomic DNA extraction was done using the DNeasy Blood and Tissue kits (Qiagen), and PCR amplification was done using the following primers *Orco*_TIDE_F1 containing the T7 promoter sequence (TAATACGACTCACTATAGGGCTCCCTTTCCCTCCCTGCAGATGA) and *Orco*_Revertant_R1 (CTGAATAAGGGCCACCCTATC). PCR product was purified using the Monarch PCR & DNA Cleanup Kit (New England BioLabs) and purified PCR products were sent for Sanger sequencing at Eurofins Genomics. CRISPR activity was analyzed using the TIDE web-based software (http://shinyapps.datacurators.nl/tide/) ^95^, by inputting the guide sequence and uploading both the test and sample sequencing files.

Eggs from G_0_ females with detectable CRISPR activities were hatched and the resulting G_1_ individuals were reared to the adult stage. G_1_ females were allowed to freely mate with *orco^16/16^* males, blood-fed and G_2_ eggs collected. G_1_ females were then sacrificed and processed as described above. Single stranded DNA oligo insertion was assessed using the Synthego (https://ice.synthego.com/#/) and TIDE (http://shinyapps.datacurators.nl/tide/) platforms. G_2_ eggs from G_1_ females with a germline integration of the donor DNA were hatched and reared to adult stage. Virgin G_2_ females were allowed to mate freely with *orco^16/16^* males. Mosquitoes were outcrossed in this manner for 3 generations. Single-pair crosses were setup among G_4_ siblings and individuals with the full length ssODN sequence insertion were chosen for homozygosing.

### Quantification and statistical analysis

Statistical analyses for all quantitative datasets from our fecundity, fertility, egg fertilization, embryonic development and sperm count assays were performed using the GraphPad Prism 9 software package (GraphPad Software, San Diego, CA, USA). All details for statistical analysis including the statistical tests used, number of trials (n), and how significance was determined can be found in the figure legends.

